# Stable nitrogen isotope analysis of amino acids as a new tool to clarify complex parasite-host interactions within marine food webs

**DOI:** 10.1101/2021.03.04.433913

**Authors:** Philip Riekenberg, Tijs Joling, Lonneke L. IJsseldijk, Andreas M. Waser, Marcel van der Meer, David W. Thieltges

## Abstract

1. Traditional bulk isotopic analysis is a pivotal tool for mapping consumer-resource interactions in food webs but has largely failed to adequately describe parasite-host relationships. Thus, parasite-host interactions remain largely understudied in food web frameworks despite these relationships increasing linkage density, connectance, and ecosystem biomass. Compound-specific stable isotopes from amino acids provides a promising novel approach that may aid in mapping parasitic interactions in food webs. However, to date it has not been applied to parasitic trophic interactions.
2. Here we use a combination of traditional bulk stable isotope analyses and compound-specific isotopic analysis of the nitrogen in amino acids to examine resource use and trophic interactions of five parasites from three hosts from a marine coastal food web (Wadden Sea, European Atlantic). By comparing isotopic compositions of bulk and amino acid nitrogen, we aimed to characterize isotopic fractionation occurring between parasites and their hosts and to clarify the trophic position of the parasites.
3. Our results showed that parasitic trophic interactions were more accurately identified when using compound-specific stable isotope analysis due to removal of underlying source isotopic variation for both parasites and hosts, and avoidance of the averaging of amino acid variability in bulk analyses through use of multiple trophic amino acids. The compound-specific method provided clear trophic discrimination factors in comparison to bulk isotope methods, however, those differences varied significantly among parasite species.
4. Amino acid compound specific isotope analysis has widely been applied to examine trophic position within food webs, but our analyses suggest that the method is particularly useful for clarifying the feeding strategies for parasitic species. Baseline isotopic information provided by source amino acids allows clear identification of the fractionation occurring due to parasite metabolism by integrating underlying isotopic variations from the host tissues. However, like for bulk isotope analysis, the application of a universal trophic discrimination factor to parasite-host relationships remains inappropriate for compound-specific stable isotope analysis. Despite this limitation, compound-specific stable isotope analysis is and will continue to be a valuable tool to increase our understanding of parasitic interactions in marine food webs.

## Introduction

Bulk tissue stable isotope analysis (hereafter δ^13^C_bulk_ or δ^15^N_bulk_ and Bulk-SIA) is routinely applied to identify resource utilization and trophic interactions within food webs (Minagawa & Wada 1984; Fry 2006). Trophic positions (TP) are identified through application of one (Post 2002; McCutchan *et al.* 2003) or several (Hussey *et al.* 2014) trophic discrimination factors (TDFs) for consumers describing the difference in isotopic composition between the consumer and their diet (Δ) for carbon or nitrogen. However, application of system-wide or group-specific TDFs do not reliably account for parasite-host relationships, which have TDFs varying from −6.7‰ to 9.0‰ (Thieltges *et al.* 2019) for δ^15^N_bulk_. These values fall well outside of the ranges typically observed for δ^15^N_bulk_ TDFs for consumers (McCutchan *et al.* 2003; Mill, Pinnegar & Polunin 2007; Caut, Angulo & Courchamp 2009) and span across parasitic species (Pinnegar, Campbell & Polunin 2001b; Dubois *et al.* 2009), feeding styles and host tissue specificity, indicating that TDFs for parasites may be specific to each relationship (*Lafferty et al.* 2008; Thieltges *et al.* 2019).

Parasitism describes a trophic interaction between two species where the parasite lives in or on the host and obtains part of, or all of, its nutritional requirements through feeding on its host, with a negative effect for the host (Lafferty & Kuris 2002). Parasitism is a widespread lifestyle, with ~40% of all organisms having at least one parasitic life-stage. Accounting for parasitic relationships, ~75% of food web interactions involve a parasite (Dobson *et al.* 2008) causing increases in linkage density (Lafferty *et al.* 2008; Sánchez Barranco *et al.* 2020), food chain length (Amundsen *et al.* 2009), and connectance (Dunne *et al.* 2013). Contributions of parasite biomass to an ecosystem can be substantial and even exceed that of top predators (Kuris *et al.* 2008; Preston *et al.* 2013). Despite widespread occurrence of parasites throughout ecosystems, these relationships have been neglected during construction of food webs since they are often incompletely understood and/or described (Marcogliese & Cone 1997; *Lafferty et al.* 2008). Incomplete characterizations of parasite-host relationships are due to the cryptic nature of interactions resulting from the difficulty of reliable sampling during the complex life-cycles of many parasite (Goater, Goater & Esch 2014).

Several mechanisms potentially contribute to variability in TDFs for parasitic relationships: 1) Mismatch between measured tissue type and tissue use by parasite, 2) Incorporation of multiple materials depending on parasite feeding style, and 3) Differing abilities in directly acquiring, anabolizing or metabolizing amino acids between parasites (*Riekenberg et al. 2021*). Many studies examining parasite-host isotopic differences used host muscle tissue despite local parasite attachment to or inhabiting another organ or tissue. Tissue mismatch can cause erroneous Δ^15^N values for parasites feeding on non-muscle host tissue (Pinnegar, Campbell & Polunin 2001a; Kamiya, Urabe & Okuda 2019). Parasite feeding styles vary, from external blood suckers to mixed diet intestinal feeders (Goater, Goater & Esch 2014) resulting in variations in parasite dietary compositions ranging from complete reliance on host tissue to predominately host gut contents (Deudero, Pinnegar & Polunin 2002; Goedknegt *et al.* 2018). Metabolic abilities of parasites such as helminths are often less complete than those found in vertebrates (Tyagi *et al.* 2015) leading to variability in biosynthesis capabilities and metabolism of compounds between species, even within the same phylum. Metabolic limitations arise when parasites can directly utilize amino acids or lipids from their hosts without requiring additional metabolism that results in variable Δ^13^C or Δ^15^N values (Deudero, Pinnegar & Polunin 2002; Power & Klein 2004; O’Grady & Dearing 2006) such as transamination or lipid synthesis pathways that occur along different metabolic pathways other than normally observed in vertebrate trophic relationships (O’Connell 2017). Impacts on Δ^15^N values resulting from unique metabolism and feeding styles can confound determination of whether a parasite is feeding on its host or supplementing with other resources (e.g. gut content) based on Bulk-SIA methods alone.

Compound-specific stable isotope analysis (CSIA) of nitrogen from amino acids (AA), hereafter δ^15^N_AA_ and AA-CSIA, provides a potential solution by simultaneously resolving baseline changes from trophic effects between parasites and hosts (Chikaraishi *et al.* 2007; Sabadel *et al.* 2016). This information comes from amino acids occurring in three types: 1) source, that undergo minimal change during metabolism and strongly retain a signal of underlying nitrogen being utilized (e.g. host or dietary N); 2) trophic, with stepwise increases as they are metabolized via transamination; and 3) metabolic, which change due to physiological processes in the animal or whether particular processing pathways are present (Popp *et al.* 2007; Chikaraishi *et al.* 2009; McMahon & McCarthy 2016). Using AAs, TPs can be estimated by using the δ^15^N value difference between a trophic and a source AA combined with system-wide values for TDFs and the Δ^15^N between trophic and source AAs in the underlying primary producers (Chikaraishi *et al.* 2009; Whiteman *et al.* 2019). In parasite-host relationships, δ^15^N_AA_ allows for isolation of the metabolism and feeding style effects because it resolves underlying shifts in baseline δ^15^N value for host or parasite to further clarify the trophic interaction. Although very promising, δ^15^N_AA_ in parasite studies has barely been utilised. A single study confirmed a large bulk Δ^15^N difference between a scarab beetle larvae and a mite (7.5‰, 2.2 TP) matched the difference found with δ^15^N_AA_ (1.8 TP) indicating a mutualistic relationship rather than parasitism between the pair (Sabadel *et al.* 2016; Sabadel, Stumbo & MacLeod 2019). To the authors knowledge there have been no studies that use δ^15^N_AA_ for parasite-host relationships with a Δ^15^N_Bulk_ value below the typically applied TDF value of 3.4‰.

In this study, we apply and compare results from both Bulk-SIA and AA-CSIA analyses applied to five parasites from three common host species from a marine coastal food web (Dutch Wadden Sea). The 5 parasite-host pairings analysed in this study are 1) the copepod *Mytilicola orientalis-*Pacific oyster (*Crassostrea gigas*), 2) the parasitic barnacle *Sacculina carcini*-shore crab (*Carcinus maenas*), 3) lung (*Pseudaliida*), 4) stomach (*Aniskis simplex*), and 5) ear nematodes *(Stenurus minor*)-harbour porpoise (*Phocoena phocoena*). We are the first to apply δ^15^N_AA_ to examine parasite-host relationships in a range of marine animals that span the breadth of trophic positions present in coastal marine food webs with oysters (*C. gigas*) as primary consumers, green shore crabs (*C. maenas*) as secondary consumers, and harbour porpoise (*P. phocoena*) as an apex predator. Here we: 1) identify whether AA-CSIA improves characterization of parasite-host relationships compared to Bulk-SIA, 2) identify the AA fractionations driving these differences, and 3) determine whether use of locally-associated host tissue improves indications of trophic relationships.

## 2) Materials and methods

Details about species included in parasite-host relationships and sample collection procedures are included in the supplemental material sections S1 and S2

### 2.1) Bulk and amino acids stable isotope analysis

All sample tissues (<1g wet weight) were frozen (−20°C) immediately after dissection and lyophilized within 2 weeks of sampling. Lyophilized tissues were ground using a mortar and pestle. Crab hepatopancreas and *S. carcini* samples were lipid extracted prior to analysis. Tissue samples were then loaded into tin capsules for Bulk-SIA analysis using a Delta V Advantage Isotope ratio mass spectrometer with a Flash 2000 Organic Element Analyzer (Thermo Fisher Scientific). The reference materials acetanilide (Biogeochemical Laboratories at Indiana University), urea, and casein (Microanalysis Ltd., Okehampton, UK) were standards for stable isotope measurements for δ^13^C and δ^15^N expressed as per mil (‰) differences from the δ^13^C value of Vienna Peedee-Belemnite Limestone (VPDB) and the δ^15^N value of atmospheric N_2_. Sample precision was ±0.1‰ and ±0.2‰ for δ^13^C and δ^15^N, respectively, for bulk materials supporting this study.

For CSIA-AA analysis, dried and homogenized tissue (2-5 mg) was lipid extracted, acid hydrolyzed, derivatized to pivaloyl- isopropyl esters and analysed for δ^15^N of AAs via gas chromatography-combustion isotope mass spectrometry using a Thermo Trace 1310 GC attached to Delta V Advantage isotope ratio mass spectrometer via an Isolink 2 following the method presented in Riekenberg et al. (2020). δ^15^N values for 14 AAs are reported: alanine (Ala), aspartic acid (Asp), glutamic acid (Glu), glycine (Gly), leucine (Leu), lysine (Lys), isoleucine (Ile), methionine (Met), phenylalanine (Phe), proline (Pro), serine (Ser), threonine (Thr), tyrosine (Tyr), and valine (Val). Precision of standards and sample amino acid δ^15^N values were <0.5‰ for 14 AAs throughout the analytical runs supporting this study. Normalization using two standard reference mixtures along with a reference spike of norleucine (Nle) added to all samples and standards which served as an internal reference calibration following the method from Yarnes and Herszage (2017).

### 2.2) Parasite-host differences and trophic position

Parasite-host differences for Δ^15^N_Bulk_ & Δ^13^C_Bulk_ were calculated for each parasite-host pair separately by subtracting the isotope ratio of host tissue from the isotope ratio of the associated parasite. For parasites of crabs and harbour porpoises this was reported for both muscle and locally-associated host tissues. Trophic position estimates were calculated using δ^15^N values from bulk material:

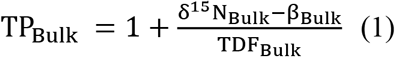

where β_Bulk_ is the δ^15^N value for primary producers in the ecosystem (9.44‰; *Goedknegt et al.* 2018), TDF_Bulk_ is the difference between consumer and their diet (3.4‰; (Minagawa & Wada 1984).

Trophic position estimates using CSIA-AA were calculated using two approaches:

1. δ^15^N values from only one trophic AA and one source AA, Glu and Phe, respectively:

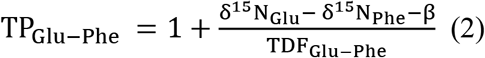

where TDF_Glu-Phe_ represents the expected stepwise increase between consumer and diet for δ^15^N value of Glu with each trophic step and β represents the difference between Glu and Phe within the primary producer at the base of the food web (Chikaraishi *et al.* 2009) A TDF_Glu-Phe_ of 6.6‰ and a β_Glu-Phe_ of 2.9‰ were applied from the meta-analysis presented in Nielsen *et al.* (2015).
2. An equation using 4 trophic and 1 source AA (TP_5AA_) as described in Nielsen *et al.* (2015):

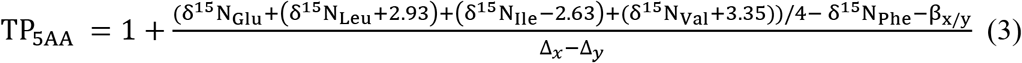

where the TP_5AA_ method incorporates more AAs than the TP_Glu-Phe_ method and is less affected by outlying measurements from individual AAs. Incorporating more source and trophic AAs reduces TP variation and gives a more precise estimation of TP. The δ^15^N_AA_ values of Leu, Ile and Val are normalized to the δ^15^N_Glu_ value using the values 2.93‰, 2.63‰, and 3.35‰, respectively (Nielsen *et al.* 2015) which allows for the calculation of a single mean δ^15^N_AA_ value using multiple trophic AAs using calculated β_x/y_ = 2.9‰ and Δ_x_−Δ_y_ = 5.9‰.

AA imbalance represents the AA concentration in host tissue compared to the parasite’s tissue composition. AA imbalance was calculated by subtracting the concentration of individual AAs (mg/g) in the locally-associated host tissue from the corresponding parasite tissue concentration (McMahon *et al.* 2015).

## 3) Results

### 3.1) Bulk isotope comparisons

Between host comparisons for bulk isotope data can be found in S3 in the supplemental material. Pairwise t-tests and Wilcoxon tests (for non-normally distributed measurements) describe δ^13^C and δ^15^N comparisons between parasites and host tissues as appropriate following Shapiro-Wilk tests for normality. *M. orientalis* (−18.5±0.3‰) and *S. carcini* (−17.7±0.6‰) had lower δ^13^C_Bulk_ values than their host muscle tissue, oyster adductor (−17.8±0.3‰; t_21_=−8.60, *p*<0.001) and crab claw-muscle (−16.7±0.8‰; t_25_=-5.26, *p*<0.001), respectively. However, the δ^13^C value of *S. carcini* did not differ from locally-associated host hepatopancreas (−17.6±0.6‰; t_25_=−1.82, *p*=0.08). The δ^13^C value of lung (−18.0±0.8‰) and stomach nematodes (−17.0±1.6‰) were similar to both harbour porpoise muscle tissue (−17.9±0.4‰; t_7_=−0.79, *p*=0.46; t_5_=1.69, *p*=0.15) and locally-associated host tissues (lung: −17.8±0.7‰; t_7_=−1.07, *p*=0.32 & stomach: −17.2±0.4‰; t_5_=0.35, *p*=0.74). The ear nematode (−16.8±0.5‰) had a higher δ ^13^C_Bulk_ value compared to harbour porpoise muscle (*p*<0.001) and locally-associated ear tissue (−19.3±1.5‰; t_6_=5.28, *p*<0.01).

*M. orientalis* (11.7±0.4‰) had no significant ^15^N_Bulk_ enrichment compared to its oyster host (11.7±0.4‰; t_21_=0.1, *p*=0.91). The δ^15^N_Bulk_ value of *S. carcini* (14.1±0.5‰) did not differ from the claw tissue of its host (14.2±0.6‰; t_25_=−1.06, *p*=0.30) but was higher compared to locally-associated hepatopancreas (13.5±0.9‰; W=58, *p*<0.01). The δ^15^N_Bulk_ value of lung nematode (17.7±1.7‰) was higher than harbour porpoise muscle tissue (15.8±1.1‰; t_7_=7.30, *p*<0.001), but comparable to locally-associated lung tissue (17.7±1.8‰; t_7_=−0.09, *p*=0.93). The δ^15^N_Bulk_ value of the stomach nematode (15.1±2.1‰) was similar to harbour porpoise muscle (15.8±1.1‰; W=6, *p*=0.43) and stomach tissue (16.6±1.3‰; W=18, *p*=0.16). The ear nematodes had the highest δ^15^N_Bulk_ value (21.8±1.0‰), higher than both harbour porpoise muscle (*p*<0.001) and locally-associated ear tissue (17.4±1.3‰; t_6_=26.55, *p*<0.001).

### 3.2) Amino acid isotope comparisons

δ^15^N_AA_ values were similar for the different tissues from the same host, with Val, Ala, Glu, Asp, Leu, Ile and Pro having higher values representative of trophic AAs, Tyr, Phe, Met, Lys and Gly, Ser, Thr having lower values associated with source and metabolic AA groupings, respectively. Pairwise t-tests (Supplemental table 1) indicated significant differences between parasite and host trophic AAs in *M.-orientalis-*oyster (Fig. 2A), *S.carcini-*crab (Fig. 2B), ear nematode-harbour porpoise (Fig. 3A) and the lung nematode-harbour porpoise (Fig. 3C) relationships, but not for the stomach nematode-harbour porpoise relationship (Fig. 3B). All parasite-host pairings were significantly different for at least one of the source AAs, but differences were variable in magnitude and direction. The largest difference observed for source AAs was for Tyr in the ear nematode-harbour porpoise pairing for muscle (18.3‰; t_5_=14.29, *p*<0.01) and ear (18.8‰; t_5_=24.25, *p*<0.01) tissues. All parasite-host pairings except for *S.carcini-*crab had significant differences in metabolic AAs that varied in magnitude and direction. Gly differences between lung nematode and locally-associated lung tissue from harbour porpoise were large and negative (−5.7‰; t_7_=−6.68, *p*<0.01) despite no difference occurring between parasite and host muscle tissue (*p*=0.7).

**Figure 2.**
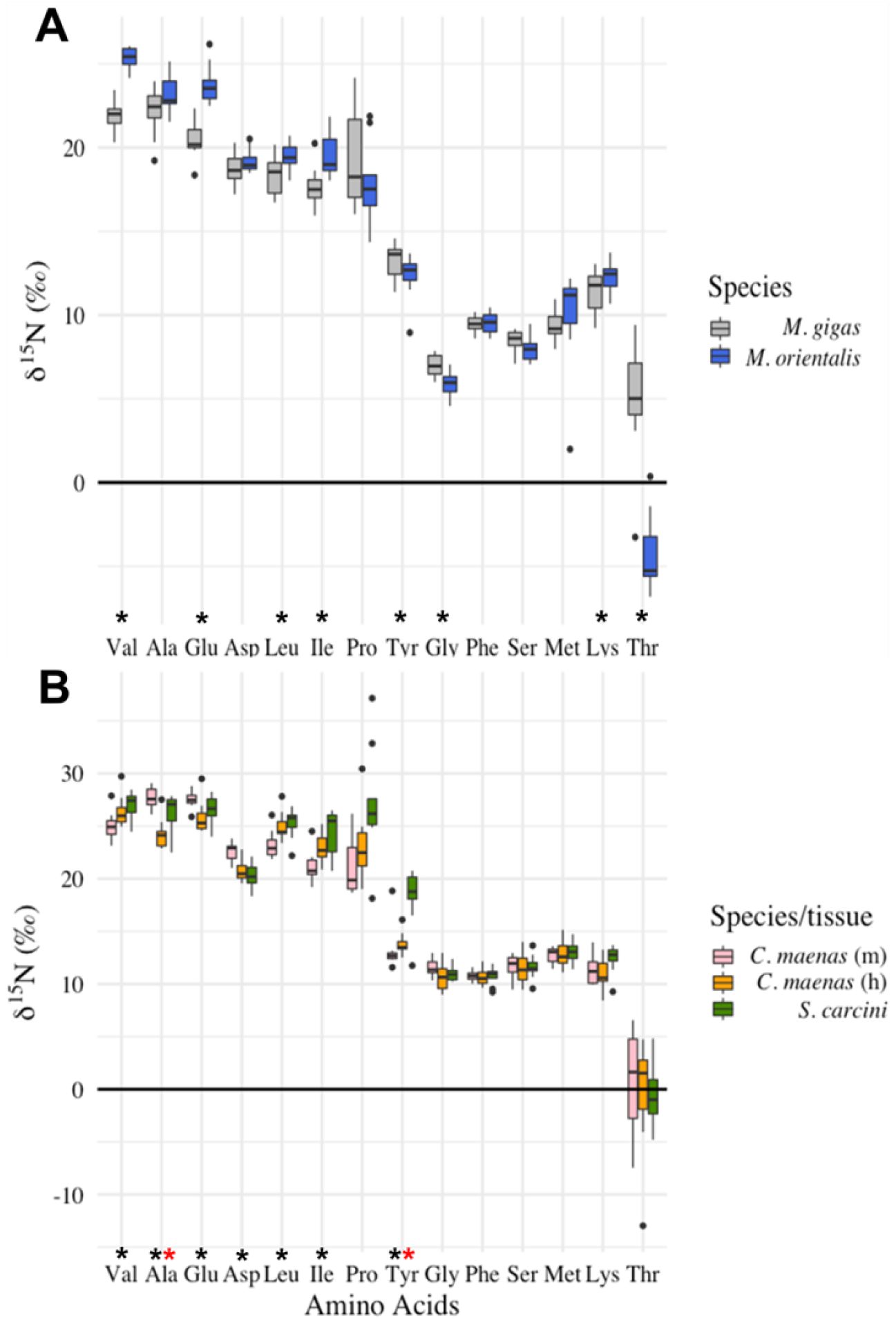
δ^15^N values for amino acids in A) *M. orientalis*-*M. gigas* and B) *S. carcini-C. maenas* parasite-host pairings. Median δ^15^N_AA_ values (solid line) are presented for 14 AAs for 12 pairings for *M. orientalis* and 11 pairings for *S. carcini*. Significant pair-wise differences (* = *p*<0.05) are indicated between parasite and host tissue using t-tests or Wilcoxon tests depending on normal distribution as indicated by Shapiro –Wilk tests for normality for each pairing. Black asterisks indicate parasite-host muscle (m) tissue differences while red asterisks indicate parasite-host local tissue differences e.g. hepatopancreas (h). Boxes and error bars represent 25th and 10th percentiles, respectively, and dots represent outliers. AA order was determined by relative fractionation difference and do not reflect the groupings defined by the PCA analysis.

**Figure 3.**
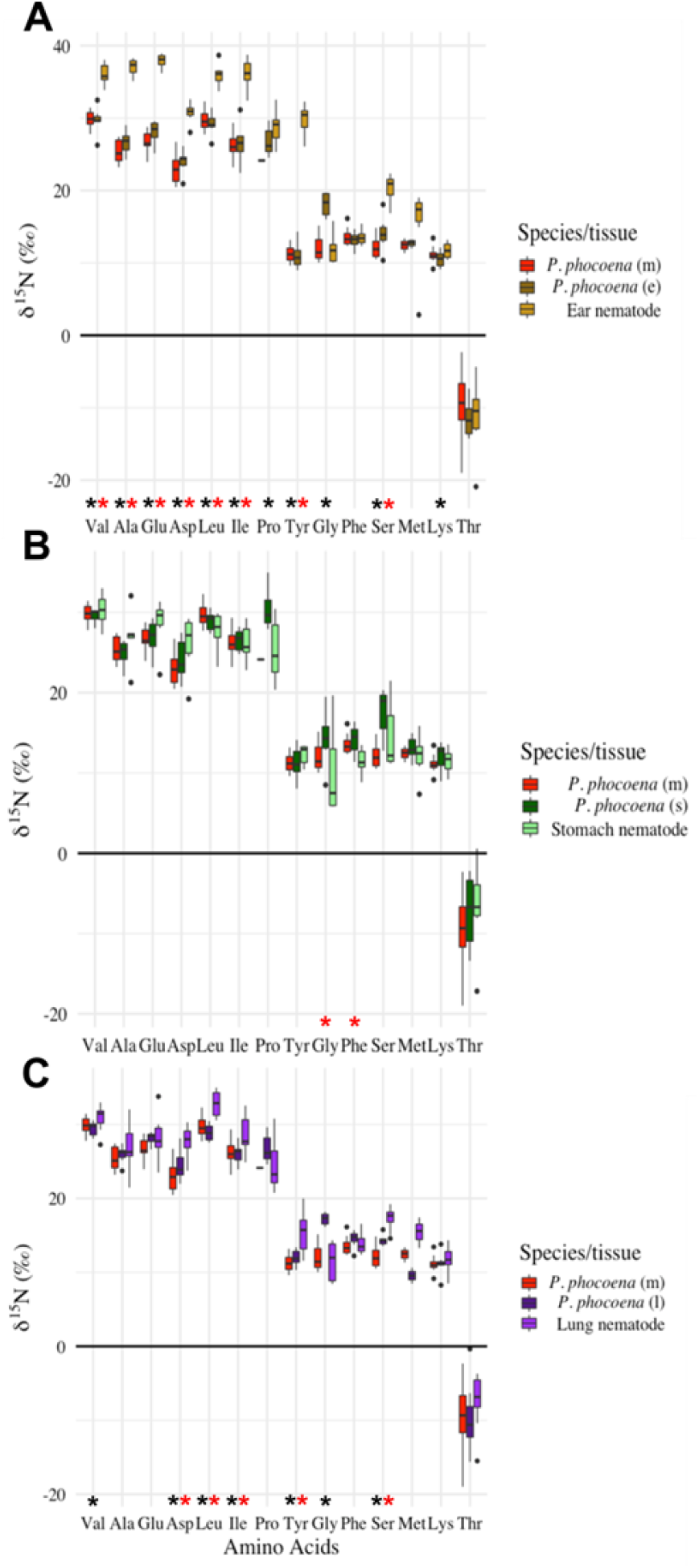
δ^15^N values for amino acids in A) ear nematodes, B) stomach nematodes, and C) lung nematodes from *P. phocoena* parasite-host pairings. Median δ^15^N_AA_ values (solid line) are presented for 14 AAs for 7 pairings for ear, 6 pairings for stomach, and 8 pairings for lung nematodes. Significant pair-wise differences (* = *p*<0.05) are indicated between parasite and host tissues using t-tests. Black asterisks indicate parasite-host muscle (m) tissue differences while red asterisks indicate parasite-host local tissue differences e.g. ear (e), stomach (s), or lung (l). Boxes and error bars represent 25th and 10th percentiles, respectively, and dots represent outliers. AA order was determined by relative fractionation difference and do not reflect the groupings defined by the PCA analysis.

**Table 1:** δ^15^N_Bulk_ & δ^13^C_Bulk_ data for each parasite and host tissue pairing.

| Host | Host tissue or parasite | No. of samples | $\delta^{13}\text{C}$ (‰) | | $\delta^{15}\text{N}$ (‰) | |
| --- | --- | --- | --- | --- | --- | --- |
|  |  |  | mean | SD | mean | SD |
| *M. giga* | Muscle | 22 | -17.8 | 0.3 | 11.7 | 0.4 |
|  | ***M. orientalis*** | 22 | -18.5 | 0.3 | 11.7 | 0.4 |
| *C. maenas* | Muscle | 26 | -16.7 | 0.8 | 14.2 | 0.6 |
|  | Hepatopancreas | 27 | -17.6 | 0.6 | 13.5 | 0.9 |
|  | ***S. carcini*** | 27 | -17.7 | 0.6 | 14.1 | 0.5 |
| *P. phocoena* | Muscle | 8 | -17.9 | 0.5 | 15.8 | 1.2 |
|  | Lung | 8 | -17.8 | 0.7 | 17.7 | 1.8 |
|  | Ear canal | 7 | -19.3 | 1.5 | 17.4 | 1.3 |
|  | Stomach | 6 | -17.2 | 0.4 | 16.6 | 1.3 |
|  | **Lung nematode** | 8 | -18 | 0.8 | 17.7 | 1.7 |
|  | **Ear nematode** | 7 | -16.8 | 0.5 | 21.8 | 1 |
|  | **Stomach nematode** | 6 | -17 | 1.6 | 15.1 | 2.1 |

Principal component analyses for δ^15^N differences between parasite-locally-associated host tissue pairs indicate that the first principal component (PC1) explained 47 to 69% and the second principal component (PC2) explained 12 to 20% of the total δ^15^N_AA_ variance (Fig. 4). The parasites *M. orientalis*, lung and ear nematodes were clearly separated from their respective hosts with non-overlapping 95% confidence intervals in multivariate space (Fig. 4A, C&D), while *S. carcini* and stomach nematodes were not separated from their host (Fig. 4 B&E.). The AAs in the principal component analysis largely grouped together according to the trophic, metabolic and source AA groupings that reflect fractionations associated with their metabolism. The ear and lung nematodes had positive loadings of similar magnitude for trophic AAs (Fig. 4 C&D.), but trophic AAs also drove contrasting negative loadings for PC1 for *M. orientalis* (Fig. 4A.). Trophic AA loadings for PC2 were near zero in ear nematodes, small for *M. orientalis* and significant (both positive and negative) for lung nematodes (Fig. 2A, C&D.). Loadings for Glu and Ala in lung nematodes were considerably different from the other trophic AAs and group with Lys, a source AA that did not differ in the lung nematode-lung tissue pairing (Fig. 4). Most loadings for source AAs were smaller across PC1, but values for PC2 remained larger in comparison to loadings observed for trophic AAs in *M. orientalis*, ear, and lung nematodes (Fig. 2A, C&D.). Source AAs were important loadings for PC2 in *M. orientalis*, ear, and lung nematodes (Fig. 2A, C&D) with ear nematode being negative versus the other two positive relationships. Thr contributed to separation across PC1 for *M. orientalis* but did not contribute considerably to separations observed for other pairings.

**Figure 4.**
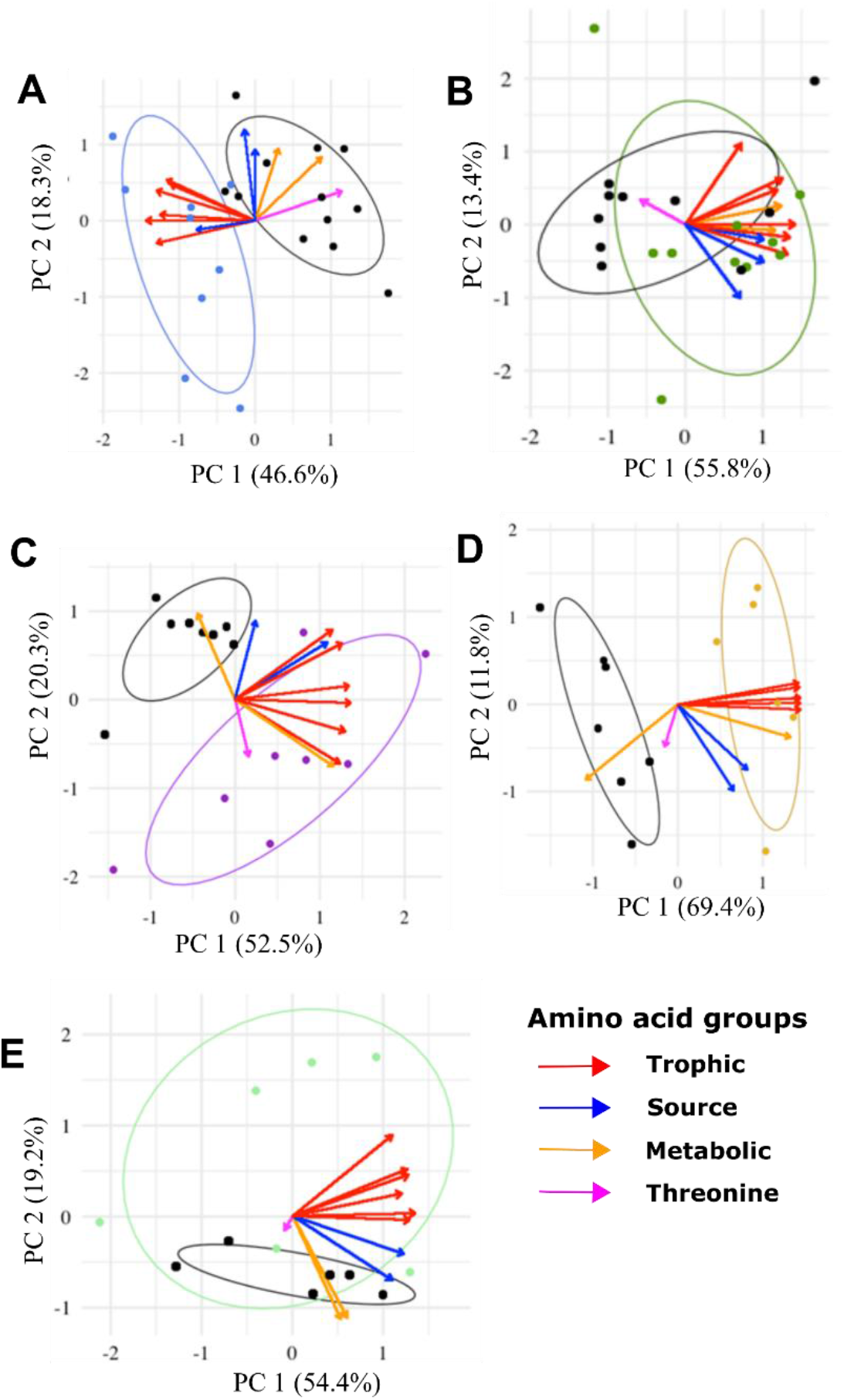
**PCA biplots of parasite-host locally-associated tissue δ^15^N_AA_ values for A)** *M. orientalis*-*M. gigas* (blue; n=9)-(black; n=12), **B)** *S. carcini*-*C. maenas* hepatopancreas tissue (green; n=10)-(black; n=11), **C)** lung nematode-*P. phocoena* lung tissue (violet; n=8)-(black; n=8)-**D)** ear nematode-*P. phocoena* ear tissue (ochre; n=6)-(black; n=7) **E)** stomach nematode-*P. phocoena* stomach tissue (light green; n=6)*-*(black; n=6). The eigenvalues among δ^15^N_AA_ values of the principal component is shown in brackets as % of variance explained. AAs are shown as eigenvectors in the plots, common fractionation behaviors are indicated by colour: red=trophic (Ala, Asp, Glu, Ile, Leu, Tyr, Val), blue=source (Phe, Lys, Met), orange= metabolic (Gly, Ser) and pink=Thr. The ellipses indicate the 95% CI for PC values of the organisms.

### 3.3) Comparing trophic position estimates between methods

TP_Bulk_ (Fig. 5A), TP_Glu-Phe_ (Fig. 5B), and TP_5AA_ (Fig. 5C) provided different trophic position estimates for parasites versus their hosts (ΔTP) between the methods (Fig. 5&6). Pairwise t-tests showed no difference in TP_Bulk_ for the *M. orientalis*-oyster pairing, but that TP_Glu-Phe_ and TP_5AA_ were significantly different (ΔTP=0.4). The *S. carcini*-crab hepatopancreas pairing had similar TPs across methods, but when compared against host (crab) muscle tissue TP_Glu-Phe_ decreased and TP_5AA_ increased. Lung parasite TP was higher than harbour porpoise muscle tissue for TP_Bulk_ and ΔTP_5AA_ but similar using TP_Glu-Phe_. When the lung nematode was compared against locally-associated lung tissue, TP_5AA_ increased, but no difference was found for ΔTP_Bulk_ or TP_Glu-Phe_. Ear nematodes had a higher TP compared to harbour porpoise muscle tissue and locally-associated ear tissue. Stomach nematodes TP_5AA_ was higher than the harbour porpoise muscle tissue, but comparable for both ΔTP_Bulk_ and TP_Glu-Phe_. However, stomach nematodes TP was higher than locally-associated stomach tissue for both TP_Glu-Phe_ and TP_5AA_, but comparable to TP_Bulk_. See supplemental table 2 for all TP pairwise t-tests for different host tissue types and Supplemental table 3 for all pairwise t-tests between TP_Bulk_ and TP_5AA_.

**Figure 5.**
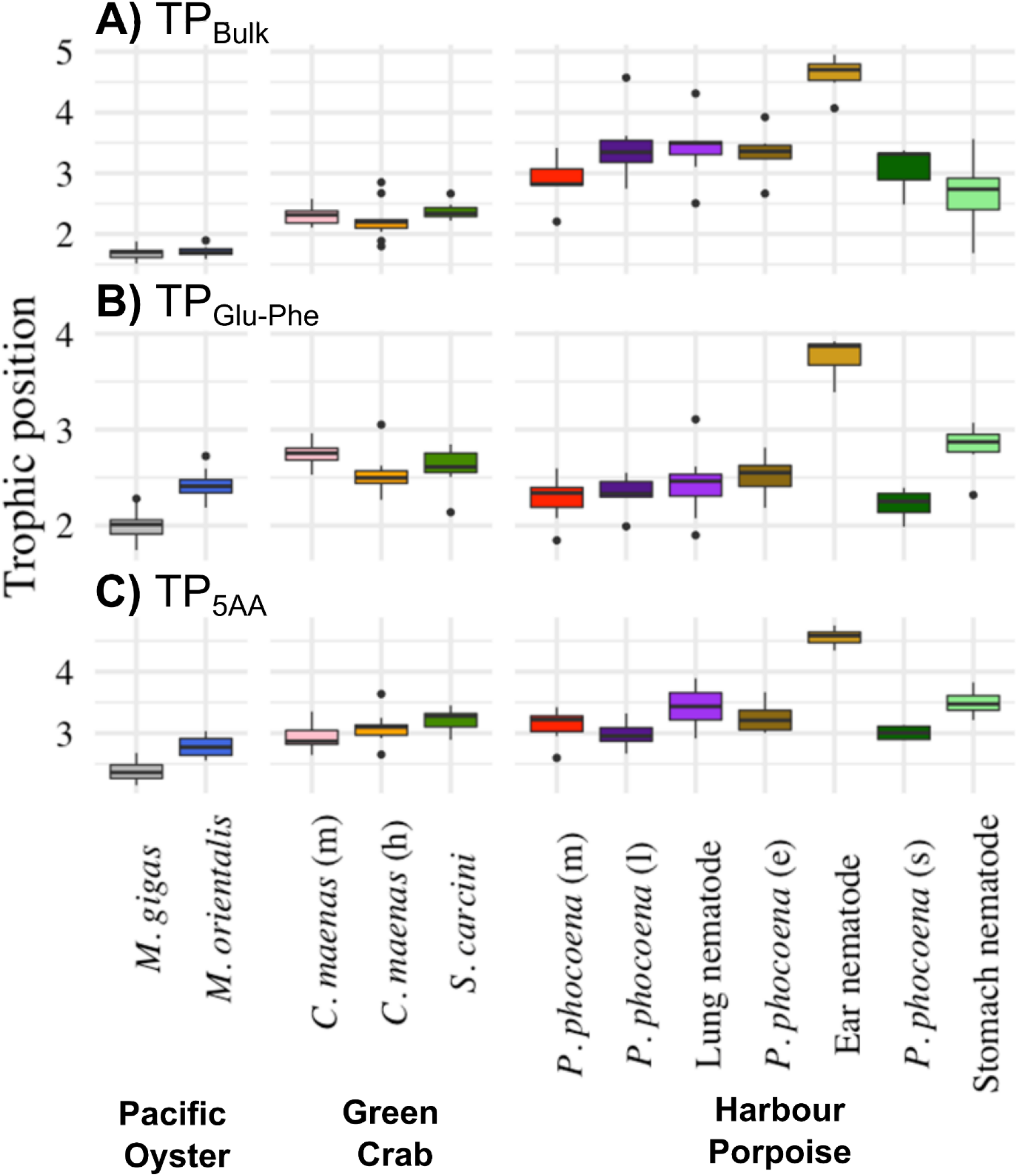
Trophic positions of parasites and hosts. Trophic positions are calculated using A) bulk δ^15^N values (TP_Bulk_), B) δ^15^N values of Glu and Phe (TP_Glu-Phe_), and C) δ^15^N values of Glu, Val, Ile, Leu and Phe (TP_5AA_). Median TP values (black line) are presented with boxes and error bars representing the 25th and 10th percentiles, respectively, and dots representing outliers. Host muscle (m) tissue or locally-associated tissues (h= hepatopancreas, l=lung, e=ear canal, s=stomach) were used for comparison of parasite attachment site.

**Figure 6:**
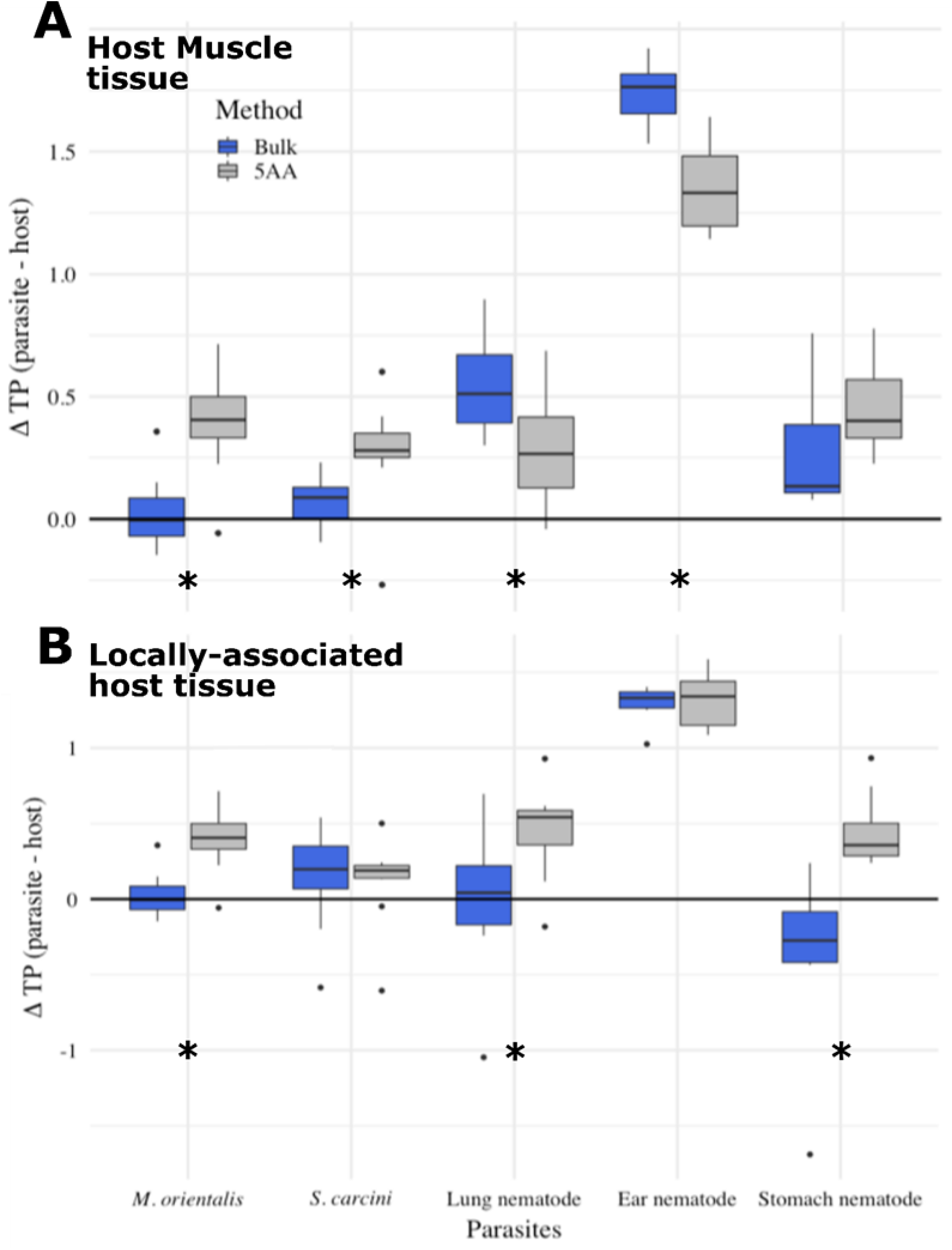
Differences in parasite-host trophic position determined by analysis of bulk nitrogen and amino acid analysis for A) parasite-host muscle tissue comparison and B) parasite-locally-associated tissue comparison. Asterisks indicate significant pairwise differences between methods at an α=0.05.

## 4) Discussion

This study identified considerable differences between Δ^15^N values determined for parasite-host pairings using Bulk-SIA (Fig.1) and AA-CSIA (Fig. 2–4) causing different TP estimates for parasites between techniques. Changes in amino acid δ^15^N values between parasite-host pairings confirmed expected groupings of source, trophic, and metabolic amino acids based on known metabolic pathways for AAs in consumers (Fig.4). However, direct uptake and utilization of AAs appears to extensively occur in the parasite (*S. carcini*) resulting in minimal fractionation and comparable TPs between the parasite and locally-associated host tissue (Fig. 4b & 6B). Differences in TP between parasite-host pairings among parasite species (Fig. 5) supports using species-specific fractionation factors when characterizing parasite relationships in food webs as well as targeted sampling of locally-associated host tissue that parasites are using. However, AA-CSIA improved the characterization of TDFs and metabolic pathways utilized for each parasitic interaction regardless of the tissue type used by providing a reliable indication of underlying δ^15^N values from the resources (e.g. tissue or dietary material) supporting each parasite.

**Figure 1:**
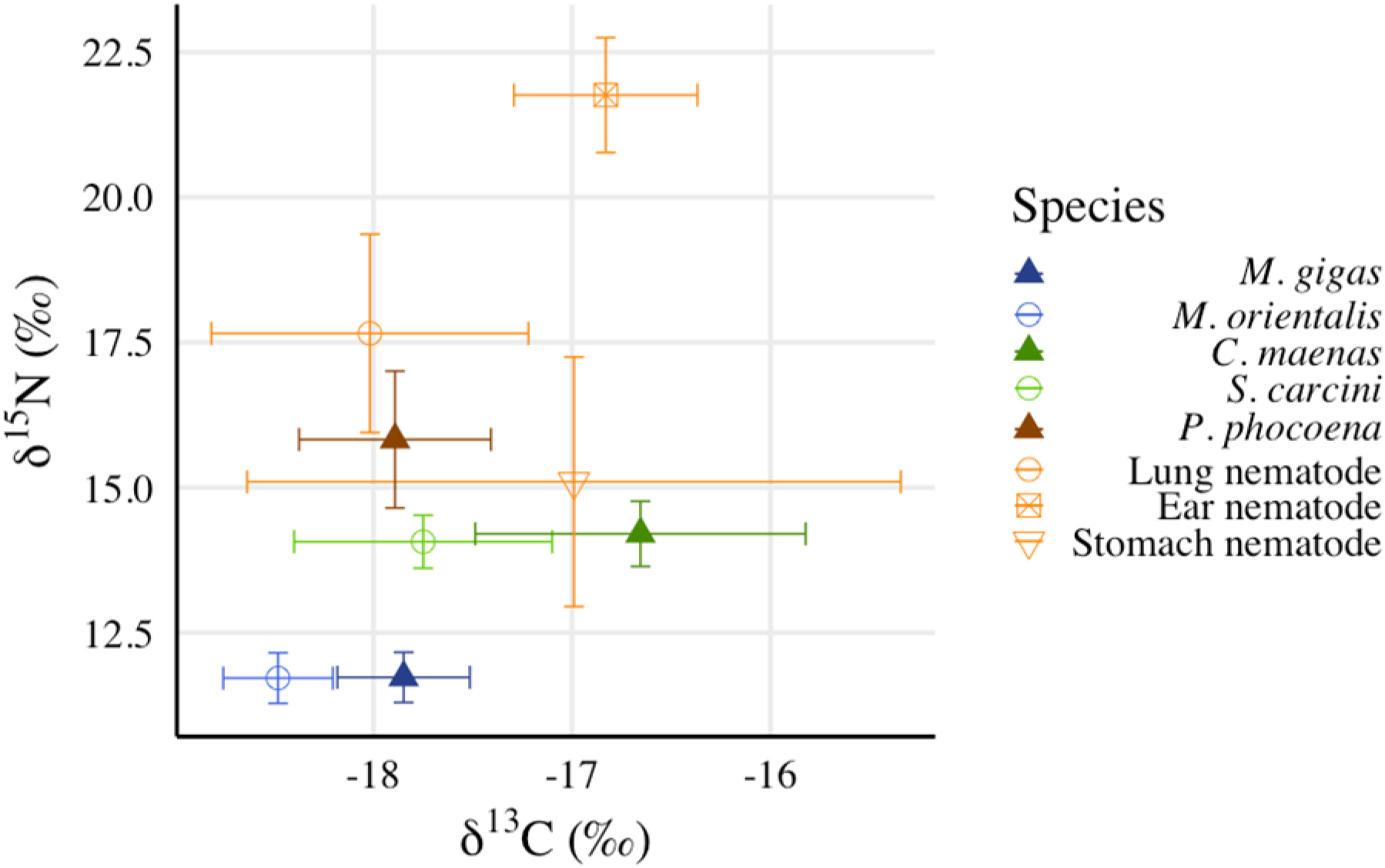
δ^15^N_Bulk_ & δ^13^C_Bulk_ data of parasites (open symbols) and hosts (full symbols). Mean δ^15^N_Bulk_ & δ^13^C_Bulk_ values for: *M. gigas* + *M. orientalis*, *C. maenas* + *S. carcini* and *P. phocoena* + parasitic nematodes (mean ± SD). For n see Table 1.

### 4.1) Fractionation differences between pairings

Δ^15^N values of parasite-host pairings varied amongst parasite feeding strategies. Both *M. orientalis* and stomach nematodes (*Aniskis simplex*) inhabit the digestive tract of their hosts (*M. gigas* & *P. phocoena*) and have access to both partially digested food and host tissue (Power & Klein 2004). AA-CSIA indicated these mixed diet feeders are both a half TP above their hosts (0.4 and 0.5, respectively; Fig. 5&6; Goedknegt *et al.* 2018). *S. carcini’s* feeding strategy is quite different from other parasites in this study as adult *S. carcini* develop extensive rootlet system to directly absorb nutrients from haemolymph associated from a variety of host tissues (Lützen 1984; Bresciani & Høeg 2001; Rowley *et al.* 2020). This direct tapping into host tissues resulted in a ΔTP_Bulk_ for *S. carcini* of ~0.1, indicating minor ^15^N fractionation during metabolism which was further confirmed by a ΔTP_5AA_ of ~0.3. Low ΔTP indicates that direct uptake of amino acids from the host is predominantly occurring with limited transamination during metabolism within *S. carcini*. These results are comparable to cestodes, who also have limited Δ^15^N values when compared to their hosts (Boag *et al.* 1998; Kamiya *et al*.2019) due to direct assimilation of compounds using a syncytial, ‘microtrich’-covered tegument (Goater *et al.* 2014) instead of digestion and metabolism of host tissue (Behrmann-Godel & Yohannes 2015). Due to direct uptake of AAs, limited AA transamination occurs prior to incorporation into parasite tissue resulting in a Δ ^15^N value that is close to zero versus the host’s diet.

Parasite feeding strategy differences do not entirely explain Δ ^15^N differences observed between parasite-host pairings. The lung (*Pseudalius inflexus* & *Torynurus convolutus*) and ear (*Stenurus minor*) nematodes were expected to have similarly high ΔTPs as both have complete digestive tracts (Goater *et al.* 2014) and feed on their locally-associated tissues of the harbour porpoise. However, the ear nematode had considerably higher ΔTP estimates for both methods (~1.3) than the lung nematode (0.3-0.5; Fig. 5 & 6) that is likely not solely a result of feeding style differences. These large variations in Δ^15^Ν values between relatively closely related parasitic species has been observed previously (Riekenberg *et al.* 2021) and are not unique to nematodes as Deudero, Pinnegar and Polunin (2002) found that different species of parasitic copepods on the same host exhibited very different δ^15^N values. Atypical ^15^N enrichments in parasites may potentially be explained by metabolic processes unique to each parasite taxon and Kamiya *et al.* (2019) and Thieltges *et al.* (2019) indicated Δ^15^Ν variations in parasite-host pairings may be related to parasite phylogenetic histories. However, despite the lung and ear parasites in this study (*P. inflexus*, *T. convolutes* & *S. minor*) all being nematodes of the same family (*Pseudaliidae)*, they maintain markedly different trophic relationships with their hosts. This result implies that utilizing the same parasite-host Δ^15^N values to determine TP_Bulk_ for closely related parasites in food webs is unlikely to provide consistently realistic results. The metabolic capabilities of nematodes appear to vary between genus to an extent where Bulk-SIA may not provide enough information to reliably resolve trophic interactions between the parasite and host without further characterization of the individual metabolic pathways and TDFs that are appropriate for each genus or species.

The considerable increase in Δ^15^Ν between the ear and lung nematodes may be explained due to relatively poor nutritional content of the ear tissue. Consumer Δ^15^N values have been observed to negatively correlate to the nutritional value of its diet (Robbins *et al*. 2010) with increased tissue C:N ratios resulting in increased TDFs due to the increased metabolic processing required to rework dietary material. The C:N ratio of ear tissue was higher than for lung tissue (4.9 vs 3.4, respectively, supplemental figure 1) and comparatively low concentrations occurred for most AAs in ear versus lung tissue. McMahon *et al.* (2015) observed a negative correlation between AA imbalance (individual trophic AA mol % in diet minus consumer, for this study individual AA concentration in host tissue minus parasite; Supplemental table 4) and TDFs for trophic AAs. In this study, Δ^15^Ν values between parasites and their host tissues were negatively correlated with AA imbalance for Leu, Lys Ser, Tyr and Val for both lung and ear nematodes (Supplemental table 5). Decreased AA availability within ear tissue is likely requiring the nematodes relying on that tissue to biosynthesise more of the AAs that are not readily available resulting in higher Δ^15^Ν_AA_ and Δ^15^Ν_Bulk_ values as more reworking occurs during metabolism. In turn, the higher availability of AAs within lung tissue results in reduced AA imbalance for the lung nematode that allows for less reworking of amino acids, more direct uptake, and smaller associated fractionations between parasite and host tissue AAs that may result in a lower Δ^15^N_AA_ (Fig. 6).

### 4.2) Changes in AAs driving fractionation

Differences in Δ^15^N values between parasites and hosts are partially driven by underlying changes occurring in individual AAs during metabolism. Examining these changes using principal component analyses revealed fractionations largely grouped along those expected for source, trophic, and metabolic AA groupings. Source AAs primarily separated across PC2 in the three pairings with significant separation between parasite and host (e.g. *M. orientalis*, lung, and ear nematodes; Fig. 4A, C&D), but variable loadings were apparent between individual source AAs (Met, Lys, Phe). This variability may indicate differences in metabolic capacity of individual parasites to metabolize source AAs or that considerable microbial reworking of source AAs occurs in the host gut content prior to parasite uptake. Given that relatively small positive fractionations are expected for source AAs during metabolism (Chikaraishi *et al.* 2009), it is surprising that source AAs occasionally have comparable loadings as trophic and metabolic AAs. Trophic AAs largely grouped together and drove separation between parasites and hosts predominately across PC1 due to the large fractionations associated with transamination during AA metabolism. The exception was Tyr, which ‘canonically’ groups within the source AAs with minimal fractionation occurring during metabolism, but consistently grouped with the trophic AAs in this analysis likely indicating that more processing occurred during metabolism which is why it has been grouped in this analysis as a trophic AA. Increased processing of Tyr indicates that potentially different metabolic processing pathways for Tyr may occur in a variety of parasites and should be examined further in future work to further characterize this relationship. The metabolic AAs Gly and Ser largely separated in the same direction as the source AAs within the parasite-host pairings, but loadings were variable across both principal components with no clear relationship between the parasite-host pairs. This variability is likely due to differing AA metabolism pathways and physiologies between the parasite species. Thr separated on its own in all of the pairings examined, which is expected due to the uniquely large negative fractionation that occurs during its metabolism (Fuller & Petzke 2017) that appears to be maintained between parasite and host, or becomes considerably more negative in δ^15^N (e.g. *M. orientalis*; Fig. 2A).

### 4.3) Host muscle versus locally-associated tissue

Large Δ^15^N values have been routinely observed between parasites and host muscle tissues and have led to the so-called ‘host-tissue isotope mismatch hypothesis’ that describes using host muscle tissue values instead of the locally-associated tissue values that the parasite is feeding upon (Pinnegar et al. 2001; Kamiya *et al.* 2019; Thieltges *et al.* 2019). By comparing both host muscle and locally-associated tissue differences in this study for all parasite-host pairings except *M. orientalis* – *M.gigas*, we found that the use of locally-associated tissue decreased the variability in ΔTP and improved agreement between the methods due to reduced fractionations associated with ΔTP_Bulk_ and ΔTP_5AA_ (Figure 6A&B). The ΔTP_AA_ analyses clearly indicate that *S. carcini* is relying on direct uptake of AAs from their host without considerable metabolic reworking and the other 4 pairings have an increasing proportion of host tissue as a portion of their diets. Increased agreement between the two methods for the *S. carcini*-crab and ear nematode-porpoise pairings is an effect of the removal of additional fractionation between locally-associated and muscle tissues. For the lung and stomach nematode-porpoise pairings, comparing against locally-associated tissue resulted in positive fractionations that indicate metabolism of N from host tissue instead of no difference or a negative relationship indicated by the bulk method (Fig. 6B). More importantly, there appears to be increased agreement between the trophic positions indicated by ΔTP_AA_ regardless of the host tissue used in the analysis (Fig. 6). This observation serves to further highlight the utility amino acid analysis that integrates variability from underlying δ^15^N baseline (e.g. biogeochemical N sources) used by the host to more accurately characterize parasite-host relationships despite the increased labour and cost associated with the method.

### 4.4) Comparing trophic position between methods

AA-CSIA enables observation of differing AA fractionations between parasite-host pairings (e.g. trophic AAs) that accounts for any variations in the underlying N supporting the parasite (e.g. source AAs). The differences between trophic position estimates described by AA-CSIA and Bulk-SIA were most apparent between *M. orientalis*-oyster (0.4 versus 0.03, respectively; Fig. 6) and the stomach nematode (*A. simplex*)-harbour porpoise (0.5 versus −0.4, respectively) for locally-associated tissues. The comparable TPs produced by Bulk-SIA for the *M. orientalis*-oyster pairing indicate that the parasite’s diet predominately consisted of digestive track contents with minimal feeding on host tissue. However, the TPs produced by AA-CSIA placed *M. orientalis* half a trophic position above *M. gigas* indicating a feeding style of a mixed gut-content / host-tissue feeder. The increase in trophic position aligns well with previous Bulk-SIA measurements taken from a *M. orientalis* pairing with an alternate bivalve host that indicated a mixed feeding style between gut content and host tissues (*M. orientalis*-*Mytilus edulis* Δ^15^N_bulk_= 1.22‰; Goedknegt *et al.* 2018). For the stomach nematode-harbour porpoise pairing, the ΔTP increased when using AA-CSIA (Fig. 6), likely driven by the strong increases in trophic AA δ^15^N values (Fig. 4E), despite decreased AA δ^15^N values for both metabolic and source AAs. AA-CSIA gives a clearer depiction of these parasite-host relationships than Bulk-SIA due to the removal of underlying N baseline variability within the wild caught hosts using source AA values (Chikaraishi *et al.* 2009) when estimating TPs. The integration of this underlying variability allows for better resolution of small differences in TP between parasite-host pairs as result of mixed feeding strategies (i.e. not solely relying on host tissue or direct uptake of substrates from the host) while accounting for any variations in the underlying biogeochemical N source values that are supporting the host within the food web. Additionally, AA-CSIA allows for the incorporation of multiple AAs which markedly reduces variability within single AA or Bulk-SIA measurements and allows a more accurate TP estimation (McCarthy *et al.* 2007; Nielsen *et al.* 2015; McMahon & McCarthy 2016; Sabadel *et al.* 2019).

### 4.5) Implications for inclusion of parasites in food webs

This work indicates that the best practice to characterize parasite-host trophic interactions is to examine locally-associated host tissue using AA-CSIA with multiple trophic and source AAs for both parasite and host materials. Use of AA-CSIA provides better clarity on parasite-host Δ^15^N values due to source AAs allowing for the integration of baseline variability for both the parasite and host that is impossible to identify with Bulk-SIA. Incorporation of multiple trophic and source AAs in the TP estimates reduces the variability that can occur within individual AAs. AA-CSIA gave a clearer indication of differences caused by the metabolism of AAs by the parasite across a broad range of trophic positions found throughout the Wadden Sea food web regardless of the type of host tissue compared against. Accounting for both parasite and host N isotopic baselines reduced uncertainty and variability in parasite-host Δ^15^_AA_ values, allowing for a more accurate analysis of these relationships regardless of host TP. Through this work, we have observed a large variability in TDFs for parasites that will likely not be adequately characterized through the application of a broad ‘universal’ TDF for parasites within food webs. Individual TDFs will likely need to be defined for each parasite species being examined, as there remain clear differences in TDFs between species contained in the same phyla (e.g. ear, lung, and stomach nematodes). This method is time intensive and costly due to the labour needed to properly derivatize the AAs prior to analysis, but AA-CSIA better characterizes the parasite-host relationships using a relatively small amount of tissue (3-5 mg). Many parasites are small, soft bodied, and difficult to cleanly sample (e.g. not easily separated from host tissue), and these limitations need to be considered prior to sample preparation. There is potential to successfully analyze smaller tissue amounts (~1 mg), but issues of host material contamination will increase as parasite sample tissue amounts decrease. Pooling samples to achieve sufficient material for analysis may be a valid option considering the additional information and clarity provided by AA-CSIA analysis on parasite-host relationships.

## Acknowledgements

We thank Jort Ossebaar and Ronald von Bommel for technical support and Ewout Adriaans, Joshua Dagoy, and Bine Keller for sampling support for this project. Funding for analysis was provided as part of Tijs Joling’s Masters internship supported by both Coastal Ocean Systems and Marine Microbiology and Biogeochemistry departments at NIOZ. Post-mortem research on harbour porpoises in the Netherlands is commissioned by the Dutch Ministry of Agriculture, Nature and Food Quality, embedded under the statutory research tasks of Wageningen UR, with project reference number WOT-04-009-045.

