## Supporting data for "Stable nitrogen isotope analysis of amino acids as a new tool to clarify complex parasite-host interactions within marine food webs"

### Abstract

1) Traditional bulk isotopic analysis is a pivotal tool for mapping consumer-resource interactions in food webs but has largely failed to adequately describe parasite-host relationships. Thus, parasite-host interactions remain largely understudied in food web frameworks despite these relationships increasing linkage density, connectance, and ecosystem biomass. Compound-specific stable isotopes from amino acids provides a promising novel approach that may aid in mapping parasitic interactions in food webs. However, to date it has not been applied to parasitic trophic interactions.

2) Here we use a combination of traditional bulk stable isotope analyses and compound-specific isotopic analysis of the nitrogen in amino acids to examine resource use and trophic interactions of five parasites from three hosts from a marine coastal food web (Wadden Sea, European Atlantic). By comparing isotopic compositions of bulk and amino acid nitrogen, we aimed to characterize isotopic fractionation occurring between parasites and their hosts and to clarify the trophic position of the parasites.

3) Our results showed that parasitic trophic interactions were more accurately identified when using compound-specific stable isotope analysis due to removal of underlying source isotopic variation for both parasites and hosts, and avoidance of the averaging of amino acid variability in bulk analyses through use of multiple trophic amino acids. The compound-specific method provided clear trophic discrimination factors in comparison to bulk isotope methods, however, those differences varied significantly among parasite species.

4) Amino acid compound specific isotope analysis has widely been applied to examine trophic position within food webs, but our analyses suggest that the method is particularly useful for clarifying the feeding strategies for parasitic species. Baseline isotopic information provided by source amino acids allows clear identification of the fractionation occurring due to parasite metabolism by integrating underlying isotopic variations from the host tissues. However, like for bulk isotope analysis, the application of a universal trophic discrimination factor to parasite-host relationships remains inappropriate for compound-specific stable isotope analysis. Despite this limitation, compound-specific stable isotope analysis is and will continue to be a valuable tool to increase our understanding of parasitic interactions in marine food webs.

**Supplemental Material**

**S1 Parasite-host relationships of the studied species**

*Crassostrea gigas* is a filter-feeding bivalve (primary consumer) that is considered an invasive species in the Wadden Sea (Troost 2010; Jung *et al.* 2019) and is host to several macroparasites including the copepod *Mytilicola orientalis* (Stock 1993; Thieltges *et al.* 2013). *M. orientalis* has a direct life cycle with a short non-feeding free-living stage, after which it permanently lives in the intestines of its host, but its feeding strategy remains incompletely characterized (Goedknegt *et al.* 2018).

*Carcinus maenas,* is a crab native to the Wadden Sea and is a secondary consumer that feeds as a generalist (Mente *et al.* 2010). *C. maenas* is host to a diverse set of parasites including nematodes, trematodes, cestodes and isopods (Zetlmeisl *et al.* 2011), but we focused on the parasitic rhizocephalan barnacle *Sacculina carcini*. Larval *S. carcini* infect their crustacean host by invading the carapace and developing during maturation an extensive root system specialised to directly absorb nutrients from a variety of tissues/internal organs (e.g. muscle, hepatopancreas and the nervous system; (Lützen 1984). *S. carcini* has no mouth, gut, respiratory organs, excretory organs or an alimentary canal during its parasitic life cycle stage (Bresciani & Høeg 2001). The effects of an infection by *S. carcini* on *C. maenas* include: parasitic castration, behavioural feminization of males, tissue changes, growth limitation or even a complete stop of the moult cycle (Powell & Rowley 2008; Kristensen *et al.* 2012; Waser *et al.* 2016). The reproductive organ of *S. carcini,* the so-called externa, develops outside of its host and is the only body part visible to the naked eye.

*Phocoena phocoena*, are small cetaceans inhabiting coastal waters and are generalist feeders on small fish like herring, sprat and anchovy (Jansen *et al.* 2013; Leopold 2015) and they are host to a diverse range of parasites including cestodes, nematodes, trematodes, crustaceans and acanthocephala (Brosens *et al.* 1996; Herreras *et al.* 1997). In this study we examined lung, ear and stomach nematodes (Goater, Goater & Esch 2014) found during necropsies of *P. phocoena* stranded on Dutch beaches. There are multiple species of nematodes parasitizing the lungs of *P. phocoena* (Lehnert, Raga & Siebert 2005) and in this study individuals from the *Pseudaliidae* family, likely *Pseudalius inflexus* or *Torynurus convolutes,* were analysed. Both nematodes have heteroxenous life cycles (Lehnert *et al.* 2010) and are very common parasites that can occur without causing health problems for their host *P. phocoena* (Lehnert, Raga & Siebert 2005; ten Doeschate *et al.* 2017), although severe infestations can limit the lung capacity, induce infections, and reduce fitness. Little is known about the life cycle of the ear nematode, *Stenurus minor*, but it is likely that invertebrates and vertebrate serve as intermediate or paratenic hosts for this parasite (Faulkner, Measures & Whoriskey 1998). Although rare, there have been reports of noise-induced hearing damage due to *S. minor* infestations in the inner ear of a harbour porpoise (Morell *et al.* 2017). The stomach nematode, *Aniskis simplex*, has a complex life cycle which includes a free-living stage, 1 or 2 intermediate crustacean hosts, and 1 or 2 paratenic fish hosts. Marine mammals are the final host of *A. simplex* (Nagasawa 1990) where the parasite occurs in the stomach and has been associated to a lowered nutritional condition of its host (ten Doeschate *et al.* 2017). It is however unclear whether *A. simplex* lowers the nutritional condition, or that harbour porpoises with a low nutritional condition are more susceptible to infections.

**S2 Sample collection**

*C. gigas* was collected by hand during low tide in April 2019 from the Mokbaai, Texel, the Netherlands (53°00'22.4"N 4°46'03.9"E). Oysters were kept alive in a tank with aerated sea water at a controlled temperature (15°C) prior to dissection. Of the 111 dissected oysters, 27 were infected with at least one *M. orientalis* individual (24% prevalence). *M. orientalis* individuals were removed from the oyster under a dissecting microscope and host tissue was sampled from the adductor muscle of the infected oysters. Parasite and host tissue samples were rinsed with deionized water after sampling, further confirmed to be free of excess tissues from host, and frozen (-20°C) immediately following dissection.

A beam trawl was used to collect *S. carcini* infected crabs in the Western Wadden Sea (52°59'50.5"N 4°51'02.4"E) in May of 2019. Infected crabs (identified by the presence of an externa) were separated from bycatch directly after each trawl on deck of the ship and uninfected *C. maenas* were returned overboard directly. A total of 27 infected *C. maenas* individuals were sampled with an estimated infection prevalence of less than 1%. Infected hosts were maintained in aerated seawater at a controlled temperature (15°C) prior to dissection. The *S. carcini* externa, *C. maenas* hepatopancreas and claw-muscle tissues were dissected under a magnifying desk lamp. The *S. carcini* and host tissues were rinsed with deionized water after sampling, further confirmed to be free of excess tissues from host, and frozen (-20°C) immediately following dissection.

Samples were collected from stranded harbour porpoises which were found dead on Dutch beaches during March 2019. Shortly after stranding, the *P. phocoena* were necropsied as part of a long-term monitoring programme at the Faculty of Veterinary Medicine, Utrecht

University, the Netherlands. Dorsal muscle (*M. Longissimus dorsi*) tissue was sampled from eight harbour porpoises and parasitic nematodes, if present, were sampled (lung, n=8; ear, n=7; stomach wall, n=5) as well as the locally associated organ tissue of the host. All samples were stored frozen (-20°C), prior to transport to NIOZ. Here, tissues were sub-sampled, rinsed with deionized water, and any remaining foreign remnant tissues from dissected tissues were removed.

The animals described in this study were free-living harbour porpoises which died of natural causes and not for the purpose of this, or other, studies. No consent from an Animal Use Committee is required, as the animals described in this study were not used for scientific or commercial testing. Consequently, animal ethics committee approval was not applicable to this work.

**S3 Bulk isotope relationships between host species**

$\delta^{13}C_{Bulk}$ values were similar between oysters and harbour porpoises (-17.8±0.3 and -17.9±0.5‰, respectively; post hoc Tukey's honest significant difference, $p>0.05$) but was significantly higher for the crabs (-16.7±0.8‰; Tukey's, $p<0.05$; One-way analysis of variance (ANOVA), $F_{2, 53}=25.18$, $p<0.001$; Fig. 1; Tab. 1.). $\delta^{15}N_{Bulk}$ values were higher for crabs and harbour porpoise (14.2±0.6 and 15.8±1.1‰; Tukey's, $p<0.05$) than for oysters (11.7±0.4‰; one-way ANOVA, $F_{2, 53} = 153.9$, $p<0.001$; Fig. 1).

**Supplemental Table 1.** Pairwise t-test results of $\delta^{15}N_{AA}$ data for all parasites and host-tissues.

| Host (tissue) | Parasite | AA | Mean | t | df | p | conf.low | conf.high |
|---|---|---|---|---|---|---|---|---|
| Oyster (o) | M. orientalis | ALA | 1.05 | 2.08 | 10 | 0.06 | -0.08 | 2.18 |
| Oyster (o) | M. orientalis | ASP | 0.42 | 1.67 | 10 | 0.13 | -0.14 | 0.98 |
| Oyster (o) | M. orientalis | GLU | 3.22 | 8.11 | 11 | <0.01 | 2.34 | 4.09 |
| Oyster (o) | M. orientalis | GLY | -1.12 | -5.88 | 11 | <0.01 | -1.53 | -0.70 |
| Oyster (o) | M. orientalis | ILE | 1.84 | 4.92 | 11 | <0.01 | 1.02 | 2.66 |
| Oyster (o) | M. orientalis | LEU | 1.14 | 4.01 | 11 | <0.01 | 0.51 | 1.76 |
| Oyster (o) | M. orientalis | LYS | 0.86 | 3.21 | 11 | <0.01 | 0.27 | 1.44 |
| Oyster (o) | M. orientalis | MET | 0.54 | 0.44 | 8 | 0.67 | -2.25 | 3.32 |
| Oyster (o) | M. orientalis | PHE | 0.04 | 0.26 | 11 | 0.8 | -0.28 | 0.35 |

| | | | | | | | | |
|---|---|---|---|---|---|---|---|---|
| Oyster (o) | M. orientalis | PRO | -1.61 | -1.45 | 6 | 0.2 | -4.32 | 1.10 |
| Oyster (o) | M. orientalis | SER | -0.44 | -1.16 | 10 | 0.27 | -1.28 | 0.40 |
| Oyster (o) | M. orientalis | THR | -8.59 | -9.75 | 9 | <0.01 | -10.58 | -6.59 |
| Oyster (o) | M. orientalis | TYR | -0.88 | -2.35 | 11 | 0.04 | -1.71 | -0.05 |
| Oyster (o) | M. orientalis | VAL | 3.41 | 11.32 | 11 | <0.01 | 2.74 | 4.07 |
| Crab (m) | S. carcini | ALA | -1.34 | -2.85 | 10 | 0.02 | -2.39 | -0.29 |
| Crab (m) | S. carcini | ASP | -2.23 | -5.12 | 10 | <0.01 | -3.20 | -1.26 |
| Crab (m) | S. carcini | GLU | -1.06 | -2.88 | 10 | 0.02 | -1.88 | -0.24 |
| Crab (m) | S. carcini | GLY | -0.54 | -2.06 | 10 | 0.07 | -1.13 | 0.05 |
| Crab (m) | S. carcini | ILE | 3.34 | 3.91 | 10 | <0.01 | 1.43 | 5.24 |
| Crab (m) | S. carcini | LEU | 2.08 | 3.21 | 10 | <0.01 | 0.64 | 3.53 |
| Crab (m) | S. carcini | LYS | 1.57 | 2.32 | 7 | 0.05 | -0.03 | 3.18 |
| Crab (m) | S. carcini | MET | 0.32 | 0.79 | 9 | 0.45 | -0.59 | 1.23 |
| Crab (m) | S. carcini | PHE | 0.00 | -0.02 | 10 | 0.99 | -0.67 | 0.66 |
| Crab (m) | S. carcini | PRO | 6.25 | 1.90 | 5 | 0.12 | -2.22 | 14.71 |
| Crab (m) | S. carcini | SER | -0.15 | -0.42 | 10 | 0.69 | -0.94 | 0.64 |
| Crab (m) | S. carcini | THR | -1.38 | -0.69 | 10 | 0.51 | -5.84 | 3.09 |
| Crab (m) | S. carcini | TYR | 4.87 | 3.16 | 8 | 0.01 | 1.32 | 8.42 |
| Crab (m) | S. carcini | VAL | 2.04 | 3.27 | 10 | <0.01 | 0.65 | 3.43 |
| Crab (h) | S. carcini | ALA | 2.21 | 3.77 | 10 | <0.01 | 0.90 | 3.51 |
| Crab (h) | S. carcini | ASP | -0.50 | -1.21 | 10 | 0.26 | -1.42 | 0.42 |
| Crab (h) | S. carcini | GLU | 0.72 | 1.23 | 10 | 0.25 | -0.58 | 2.03 |
| Crab (h) | S. carcini | GLY | 0.42 | 1.14 | 10 | 0.28 | -0.40 | 1.25 |
| Crab (h) | S. carcini | ILE | 1.60 | 2.10 | 10 | 0.06 | -0.10 | 3.29 |
| Crab (h) | S. carcini | LEU | 0.32 | 0.61 | 10 | 0.56 | -0.86 | 1.51 |
| Crab (h) | S. carcini | LYS | 1.45 | 1.93 | 9 | 0.09 | -0.25 | 3.15 |
| Crab (h) | S. carcini | MET | 0.32 | 0.76 | 9 | 0.46 | -0.63 | 1.27 |
| Crab (h) | S. carcini | PHE | 0.09 | 0.38 | 10 | 0.71 | -0.46 | 0.65 |
| Crab (h) | S. carcini | PRO | 1.23 | 0.61 | 5 | 0.57 | -3.91 | 6.37 |
| Crab (h) | S. carcini | SER | 0.15 | 0.40 | 10 | 0.7 | -0.69 | 0.99 |
| Crab (h) | S. carcini | THR | -0.36 | -0.17 | 10 | 0.87 | -4.95 | 4.23 |
| Crab (h) | S. carcini | TYR | 4.56 | 5.90 | 9 | <0.01 | 2.81 | 6.31 |
| Crab (h) | S. carcini | VAL | 0.68 | 1.27 | 10 | 0.23 | -0.52 | 1.88 |
| HP (m) | Lung nematode | ALA | 1.56 | 1.75 | 7 | 0.12 | -0.54 | 3.67 |
| HP (m) | Lung nematode | ASP | 4.67 | 7.86 | 7 | <0.01 | 3.27 | 6.08 |
| HP (m) | Lung nematode | GLU | 1.47 | 1.93 | 7 | 0.09 | -0.33 | 3.26 |
| HP (m) | Lung nematode | GLY | -0.52 | -0.43 | 7 | 0.68 | -3.42 | 2.37 |
| HP (m) | Lung nematode | ILE | 2.35 | 6.23 | 7 | <0.01 | 1.46 | 3.24 |
| HP (m) | Lung nematode | LEU | 3.00 | 9.00 | 7 | <0.01 | 2.21 | 3.79 |
| HP (m) | Lung nematode | LYS | 0.54 | 1.82 | 7 | 0.11 | -0.16 | 1.25 |
| HP (m) | Lung nematode | PHE | 0.26 | 0.62 | 7 | 0.55 | -0.73 | 1.26 |
| HP (m) | Lung nematode | SER | 5.09 | 5.40 | 7 | <0.01 | 2.86 | 7.32 |
| HP (m) | Lung nematode | THR | 2.28 | 1.55 | 7 | 0.16 | -1.19 | 5.75 |

| | | | | | | | | |
|---|---|---|---|---|---|---|---|---|
| HP (m) | Lung nematode | TYR | 4.13 | 5.12 | 6 | <0.01 | 2.16 | 6.10 |
| HP (m) | Lung nematode | VAL | 1.05 | 2.27 | 7 | 0.06 | -0.04 | 2.14 |
| HP (lu) | Lung nematode | ALA | 1.00 | 1.03 | 7 | 0.34 | -1.29 | 3.29 |
| HP (lu) | Lung nematode | ASP | 3.41 | 5.26 | 7 | <0.01 | 1.88 | 4.94 |
| HP (lu) | Lung nematode | GLU | 0.12 | 0.11 | 7 | 0.91 | -2.26 | 2.49 |
| HP (lu) | Lung nematode | GLY | -5.73 | -6.68 | 7 | <0.01 | -7.76 | -3.70 |
| HP (lu) | Lung nematode | ILE | 2.56 | 4.00 | 7 | <0.01 | 1.05 | 4.07 |
| HP (lu) | Lung nematode | LEU | 3.90 | 11.38 | 7 | <0.01 | 3.09 | 4.71 |
| HP (lu) | Lung nematode | LYS | 0.51 | 2.04 | 7 | 0.08 | -0.08 | 1.10 |
| HP (lu) | Lung nematode | PHE | -0.64 | -1.30 | 7 | 0.23 | -1.79 | 0.52 |
| HP (lu) | Lung nematode | SER | 2.97 | 5.03 | 7 | <0.01 | 1.57 | 4.36 |
| HP (lu) | Lung nematode | THR | 2.34 | 1.59 | 7 | 0.16 | -1.15 | 5.82 |
| HP (lu) | Lung nematode | TYR | 3.73 | 4.45 | 5 | <0.01 | 1.57 | 5.88 |
| HP (lu) | Lung nematode | VAL | 1.43 | 3.72 | 7 | <0.01 | 0.52 | 2.34 |
| HP (m) | Ear nematode | ALA | 11.48 | 20.43 | 6 | <0.01 | 10.11 | 12.86 |
| HP (m) | Ear nematode | ASP | 7.41 | 10.17 | 6 | <0.01 | 5.62 | 9.19 |
| HP (m) | Ear nematode | GLU | 11.04 | 18.21 | 6 | <0.01 | 9.55 | 12.52 |
| HP (m) | Ear nematode | GLY | 0.30 | 0.43 | 6 | 0.68 | -1.41 | 2.01 |
| HP (m) | Ear nematode | ILE | 9.93 | 10.79 | 6 | <0.01 | 7.68 | 12.18 |
| HP (m) | Ear nematode | LEU | 5.86 | 8.86 | 6 | <0.01 | 4.24 | 7.48 |
| HP (m) | Ear nematode | LYS | 0.57 | 1.07 | 6 | 0.32 | -0.73 | 1.86 |
| HP (m) | Ear nematode | MET | 4.99 | 5.01 | 1 | 0.13 | -7.66 | 17.63 |
| HP (m) | Ear nematode | PHE | 0.28 | 0.66 | 6 | 0.53 | -0.76 | 1.33 |
| HP (m) | Ear nematode | SER | 8.45 | 9.37 | 6 | <0.01 | 6.24 | 10.65 |
| HP (m) | Ear nematode | THR | -2.19 | -1.07 | 5 | 0.33 | -7.44 | 3.06 |
| HP (m) | Ear nematode | TYR | 18.30 | 14.29 | 5 | <0.01 | 15.00 | 21.59 |
| HP (m) | Ear nematode | VAL | 6.25 | 14.49 | 6 | <0.01 | 5.19 | 7.30 |
| HP (ec) | Ear nematode | ALA | 10.40 | 17.84 | 6 | <0.01 | 8.97 | 11.82 |
| HP (ec) | Ear nematode | ASP | 6.78 | 9.78 | 6 | <0.01 | 5.09 | 8.48 |
| HP (ec) | Ear nematode | GLU | 9.83 | 17.29 | 6 | <0.01 | 8.44 | 11.22 |
| HP (ec) | Ear nematode | GLY | -6.19 | -7.13 | 6 | <0.01 | -8.32 | -4.07 |
| HP (ec) | Ear nematode | ILE | 9.58 | 8.72 | 6 | <0.01 | 6.89 | 12.27 |
| HP (ec) | Ear nematode | LEU | 6.75 | 10.89 | 6 | <0.01 | 5.24 | 8.27 |
| HP (ec) | Ear nematode | LYS | 1.20 | 3.34 | 6 | 0.02 | 0.32 | 2.09 |
| HP (ec) | Ear nematode | MET | 1.74 | 0.45 | 3 | 0.68 | -10.42 | 13.89 |
| HP (ec) | Ear nematode | PHE | 0.40 | 1.19 | 6 | 0.28 | -0.42 | 1.22 |
| HP (ec) | Ear nematode | SER | 6.23 | 6.44 | 6 | <0.01 | 3.86 | 8.59 |
| HP (ec) | Ear nematode | THR | -0.27 | -0.09 | 5 | 0.93 | -7.83 | 7.29 |
| HP (ec) | Ear nematode | TYR | 18.82 | 24.25 | 5 | <0.01 | 16.82 | 20.82 |
| HP (ec) | Ear nematode | VAL | 6.42 | 9.87 | 6 | <0.01 | 4.82 | 8.01 |
| HP (m) | Stomach nematode | ALA | 2.31 | 1.20 | 5 | 0.28 | -2.63 | 7.25 |
| HP (m) | Stomach nematode | ASP | 3.20 | 1.45 | 5 | 0.21 | -2.48 | 8.88 |
| HP (m) | Stomach nematode | GLU | 2.59 | 1.40 | 5 | 0.22 | -2.17 | 7.36 |

| | | | | | | | | |
|---|---|---|---|---|---|---|---|---|
| HP (m) | Stomach nematode | GLY | -1.05 | -0.42 | 5 | 0.69 | -7.42 | 5.32 |
| HP (m) | Stomach nematode | ILE | 0.85 | 0.65 | 5 | 0.54 | -2.50 | 4.20 |
| HP (m) | Stomach nematode | LEU | -1.81 | -1.29 | 5 | 0.25 | -5.42 | 1.81 |
| HP (m) | Stomach nematode | LYS | 1.25 | 2.00 | 5 | 0.1 | -0.35 | 2.85 |
| HP (m) | Stomach nematode | MET | -2.93 | -1.31 | 1 | 0.42 | -31.39 | 25.53 |
| HP (m) | Stomach nematode | PHE | -1.36 | -1.60 | 5 | 0.17 | -3.54 | 0.82 |
| HP (m) | Stomach nematode | SER | 3.00 | 1.61 | 5 | 0.17 | -1.79 | 7.79 |
| HP (m) | Stomach nematode | THR | 2.69 | 1.02 | 5 | 0.35 | -4.07 | 9.44 |
| HP (m) | Stomach nematode | TYR | 1.39 | 2.27 | 5 | 0.07 | -0.18 | 2.96 |
| HP (m) | Stomach nematode | VAL | 1.16 | 1.69 | 5 | 0.15 | -0.60 | 2.91 |
| HP (s) | Stomach nematode | ALA | 2.08 | 1.41 | 5 | 0.22 | -1.71 | 5.86 |
| HP (s) | Stomach nematode | ASP | 1.93 | 1.22 | 5 | 0.28 | -2.14 | 6.01 |
| HP (s) | Stomach nematode | GLU | 1.72 | 1.31 | 5 | 0.25 | -1.65 | 5.10 |
| HP (s) | Stomach nematode | GLY | -4.14 | -2.80 | 5 | 0.04 | -7.94 | -0.34 |
| HP (s) | Stomach nematode | ILE | -0.39 | -0.41 | 5 | 0.7 | -2.81 | 2.04 |
| HP (s) | Stomach nematode | LEU | -1.29 | -1.22 | 5 | 0.28 | -4.01 | 1.42 |
| HP (s) | Stomach nematode | LYS | -0.50 | -0.82 | 5 | 0.45 | -2.05 | 1.05 |
| HP (s) | Stomach nematode | MET | -0.99 | -0.99 | 5 | 0.37 | -3.56 | 1.58 |
| HP (s) | Stomach nematode | PHE | -2.71 | -2.59 | 5 | 0.05 | -5.40 | -0.02 |
| HP (s) | Stomach nematode | PRO | -4.00 | -1.48 | 3 | 0.23 | -12.59 | 4.59 |
| HP (s) | Stomach nematode | SER | -3.05 | -1.01 | 5 | 0.36 | -10.80 | 4.70 |
| HP (s) | Stomach nematode | THR | 0.41 | 0.14 | 5 | 0.89 | -7.20 | 8.03 |
| HP (s) | Stomach nematode | TYR | 1.07 | 1.16 | 5 | 0.3 | -1.31 | 3.44 |
| HP (s) | Stomach nematode | VAL | 0.70 | 0.84 | 5 | 0.44 | -1.45 | 2.86 |

**Supplemental Table 2.** Pairwise t-test of the 3 TP calculations for all parasites and host-tissues.

| Host (tissue) | Parasite | TPmethode | Mean | t | df | p | conf.low | conf.high |
|---|---|---|---|---|---|---|---|---|
| Oyster (o) | M. orientalis | Bulk | 0.03 | 0.63 | 11 | 0.54 | -0.06 | 0.11 |
| Oyster (o) | M. orientalis | Glu-Phe | 0.42 | 7.53 | 11 | <0.01 | 0.30 | 0.54 |
| Oyster (o) | M. orientalis | 5AA | 0.40 | 6.93 | 11 | <0.01 | 0.27 | 0.53 |
| Crab (m) | S. carcini | Bulk | 0.07 | 2.19 | 10 | 0.05 | 0.00 | 0.14 |
| Crab (m) | S. carcini | Glu-Phe | -0.14 | -3.04 | 10 | 0.01 | -0.24 | -0.04 |
| Crab (m) | S. carcini | 5AA | 0.27 | 4.30 | 10 | <0.01 | 0.13 | 0.41 |
| Crab (h) | S. carcini | Bulk | 0.15 | 1.55 | 10 | 0.15 | -0.06 | 0.36 |
| Crab (h) | S. carcini | Glu-Phe | 0.08 | 1.07 | 10 | 0.31 | -0.09 | 0.25 |
| Crab (h) | S. carcini | 5AA | 0.12 | 1.51 | 10 | 0.16 | -0.06 | 0.31 |
| HP (m) | Lung nematode | Bulk | 0.54 | 7.31 | 7 | <0.01 | 0.36 | 0.71 |
| HP (m) | Lung nematode | Glu-Phe | 0.16 | 1.23 | 7 | 0.26 | -0.15 | 0.46 |
| HP (m) | Lung nematode | 5AA | 0.29 | 2.98 | 7 | 0.02 | 0.06 | 0.52 |
| HP (lu) | Lung nematode | Bulk | -0.02 | -0.09 | 7 | 0.93 | -0.44 | 0.41 |
| HP (lu) | Lung nematode | Glu-Phe | 0.10 | 0.66 | 7 | 0.53 | -0.25 | 0.45 |
| HP (lu) | Lung nematode | 5AA | 0.45 | 3.74 | 7 | <0.01 | 0.16 | 0.73 |
| HP (m) | Ear nematode | Bulk | 1.74 | 34.31 | 6 | <0.01 | 1.61 | 1.86 |
| HP (m) | Ear nematode | Glu-Phe | 1.41 | 14.32 | 6 | <0.01 | 1.17 | 1.66 |
| HP (m) | Ear nematode | 5AA | 1.35 | 18.87 | 6 | <0.01 | 1.18 | 1.53 |
| HP (ec) | Ear nematode | Bulk | 1.29 | 26.55 | 6 | <0.01 | 1.17 | 1.41 |
| HP (ec) | Ear nematode | Glu-Phe | 1.24 | 21.11 | 6 | <0.01 | 1.10 | 1.38 |
| HP (ec) | Ear nematode | 5AA | 1.31 | 17.99 | 6 | <0.01 | 1.13 | 1.49 |
| HP (m) | Stomach nematode | Bulk | 0.01 | 0.05 | 5 | 0.96 | -0.75 | 0.78 |
| HP (m) | Stomach nematode | Glu-Phe | 0.52 | 3.00 | 5 | 0.03 | 0.08 | 0.96 |
| HP (m) | Stomach nematode | 5AA | 0.35 | 2.55 | 5 | 0.05 | 0.00 | 0.70 |
| HP (s) | Stomach nematode | Bulk | -0.43 | -1.59 | 5 | 0.17 | -1.12 | 0.26 |
| HP (s) | Stomach nematode | Glu-Phe | 0.58 | 6.42 | 5 | <0.01 | 0.35 | 0.82 |
| HP (s) | Stomach nematode | 5AA | 0.49 | 4.27 | 5 | <0.01 | 0.20 | 0.79 |

**Supplemental Table 3.** Pairwise tests of the difference between the Bulk and 5AA Delta (parasite-host) TP calculations.

| Host (tissue) | Parasite | Test | Mean | t / w | df | p | conf.low | conf.high |
|---|---|---|---|---|---|---|---|---|
| Oyster (o) | M. orientalis | Paired t-test | -0.375 | -10.87 | 11 | <0.01 | -0.451 | -0.299 |
| Crab (m) | S. carcini | Wilcoxon | -0.228 | 10 | | 0.045 | -0.353 | -0.065 |
| HP (m) | Lung nematode | Paired t-test | 0.249 | 3.60 | 7 | <0.01 | 0.085 | 0.413 |
| HP (m) | Ear nematode | Paired t-test | 0.383 | 7.91 | 6 | <0.01 | 0.265 | 0.502 |
| HP (m) | Stomach nematode | Paired t-test | -0.168 | -0.81 | 4 | 0.465 | -0.747 | 0.411 |
| Crab (h) | S. carcini | Wilcoxon | -0.004 | 32 | | 0.965 | -0.142 | 0.171 |
| HP (lu) | Lung nematode | Paired t-test | -0.304 | -3.04 | 6 | 0.023 | -0.548 | -0.059 |
| HP (ec) | Ear nematode | Paired t-test | -0.024 | -0.27 | 6 | 0.793 | -0.234 | 0.187 |
| HP (s) | Stomach nematode | Paired t-test | -1.092 | -3.81 | 4 | 0.019 | -1.888 | -0.297 |

**Supplemental Table 4.** AA imbalance (individual AA concentration in host minus parasite) for all five parasites, Mean ± SD.

| AA | *M. orienetalis* | *S. carcini* | Lung nem. | Ear nem. | Stomach nem. |
|---|---|---|---|---|---|
| **ALA** | 2.29 ± 1.02 | -0.44 ± 1.01 | 2.25 ± 1.36 | 0.92 ± 2.07 | 0.09 ± 1.88 |
| **ASP** | 1.73 ± 1.46 | -0.31 ± 1.08 | 0.57 ± 1.23 | -1.85 ± 1.16 | -0.71 ± 1.69 |
| **GLU** | 3.14 ± 1.52 | -1.25 ± 1.28 | -0.09 ± 1.39 | -2.55 ± 1.73 | 0.51 ± 1.38 |
| **GLY** | 1.28 ± 1.09 | 0.01 ± 1.65 | 3.44 ± 3.72 | 3.97 ± 6.15 | 2.36 ± 2.33 |
| **ILE** | 1.35 ± 0.66 | -1.06 ± 1.67 | -0.01 ± 1.56 | -0.7 ± 0.26 | 0.3 ± 0.75 |
| **LEU** | 1.61 ± 0.83 | -0.78 ± 0.96 | 1.79 ± 0.96 | -1.02 ± 0.63 | 1.79 ± 1.22 |
| **LYS** | 0.45 ± 1.67 | -1.17 ± 1.06 | 0.19 ± 0.95 | -1.5 ± 1.39 | -3.05 ± 1.5 |
| **MET** | 0.56 ± 0.42 | -0.16 ± 0.25 | -0.14 ± 0.35 | -0.12 ± 0.22 | -0.09 ± 0.31 |
| **PHE** | 0.34 ± 0.37 | -0.01 ± 0.43 | 0.5 ± 0.73 | -0.75 ± 0.7 | 0.38 ± 0.51 |
| **PRO** | -1.88 ± 0.71 | -0.49 ± 0.56 | 0.3 ± 0.74 | -0.39 ± 0.99 | -1.44 ± 1.92 |
| **SER** | 0.08 ± 0.34 | -0.81 ± 0.57 | 0.42 ± 0.3 | -0.72 ± 0.49 | 0.64 ± 0.6 |
| **THR** | 0.17 ± 0.31 | -0.22 ± 0.52 | 0.12 ± 0.54 | -0.87 ± 0.95 | -0.86 ± 1.02 |
| **TYR** | 0.26 ± 0.3 | -0.53 ± 0.37 | 0.22 ± 0.31 | -0.65 ± 0.41 | 0.04 ± 0.32 |
| **VAL** | 1.94 ± 1.02 | -0.84 ± 0.9 | 1.52 ± 0.8 | -0.19 ± 0.72 | 0.4 ± 0.83 |

**Supplemental Table 5.** Pearson correlation coefficients of $\mathbf{\Delta^{15}N_{AA}}$ with **C:N** ratio of associated host tissue, $\mathbf{\Delta^{15}N_{Bulk}}$ and **AA imbalance** (of AA in question) for the lung (n=8) and ear nematode (n=6) of *P. phocoena*. A p-value of <0.05 is indicated as * and <0.01 as **.

| $\Delta^{15}N_{AA}$ | C:N | $\Delta^{15}N_{Bulk}$ | AA-Imbalance |
|---|---|---|---|
| **ALA** | 0.61* | 0.91** | -0.26 |
| **ASP** | 0.71** | 0.91** | -0.45 |
| **GLU** | 0.64* | 0.94** | -0.47 |
| **GLY** | 0.15 | -0.1 | 0.35 |
| **ILE** | 0.69** | 0.9** | -0.2 |
| **LEU** | 0.69** | 0.68** | -0.69** |
| **LYS** | 0.55* | 0.54* | -0.18 |
| **MET** | -0.98* | 0.97* | -0.77 |
| **PHE** | 0.38 | 0.57* | -0.65* |
| **PRO** | 0.83** | 0.64* | 0.16 |
| **SER** | 0.54* | 0.59* | -0.66* |
| **THR** | -0.74** | -0.54 | -0.1 |
| **TYR** | 0.55 | 0.95** | -0.74** |
| **VAL** | 0.71** | 0.87** | -0.65* |

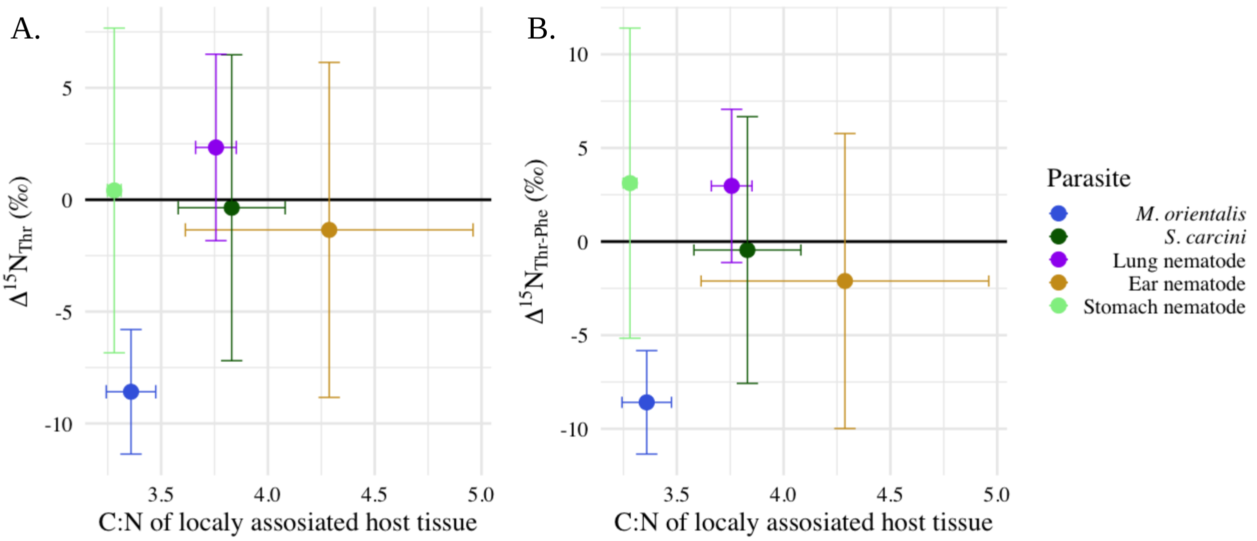


**Supplemental Figure 1. Effect of food quality (C:N) on $\Delta^{15}N_{Thr}$.** C:N ratio of locally associated host tissue for *M. orientalis* (n=9), *S. carcini* (n=11), lung- (n=8), ear- (n=6) and stomach (n=6) nematodes. **A)** $\Delta^{15}N_{Thr}$ and **B)** $\Delta^{15}N_{Thr\text{-}Phe}$. Mean ± SD are displayed. There is no linear correlation between C:N ratio and $\Delta^{15}N_{Thr}$ or $\Delta^{15}N_{Thr\text{-}Phe}$.
